# Proteome thermal stability reflects organ physiology and identifies drug-target engagement in vivo

**DOI:** 10.1101/500306

**Authors:** Jessica Perrin, Thilo Werner, Nils Kurzawa, Dorothee D. Childs, Mathias Kalxdorf, Anna Rutkowska, Daniel Poeckel, Daniel C. Sevin, Eugenia Stonehouse, Katrin Strohmer, Bianca Heller, Douglas W. Thomson, Johanna Vappiani, Jana Krause, H. Christian Eberl, Christina E. Rau, Holger Franken, Wolfgang Huber, Maria Faelth-Savitski, Mikhail M. Savitski, Marcus Bantscheff, Giovanna Bergamini

## Abstract

Studying biological processes at a molecular level and monitoring drug-target interactions is established for simple cell systems but challenging in vivo. We introduce and apply a methodology for proteome-wide thermal stability measurements to characterize organ physiology and activity of many fundamental biological processes across tissues, such as energy metabolism and protein homeostasis. This method, termed tissue thermal proteome profiling (tissue-TPP), also enabled target and off-target identification and occupancy measurements in tissues derived from animals dosed with the non-covalent histone deacetylase inhibitor, panobinostat. Finally, we devised blood-CETSA, a thermal stability-based method to monitor target engagement in whole blood. Our study generates the first proteome-wide map of protein thermal stability in tissue and provides tools that will be of great impact for translational research.

## Main Text

Thermal proteome profiling (TPP) determines thermal stability across the proteome by measuring the soluble fractions of proteins upon heating cells and lysates to a range of temperatures. It also enables the proteome-wide assessment of drug-protein interactions by combining quantitative mass spectrometry (*1*) with the cellular thermal shift assay, (CETSA)(*2*)(*3*)(*4*).

Beyond the identification of direct drug targets (*4*) TPP detects drug treatment induced perturbations of metabolic and signaling pathways (*5*)(*3*)(*6*) and modulation of protein complexes regulating complex biological processes (*7*)(*8*)(*9*) in *in vitro* cellular systems. The CETSA methodology has recently been applied to tissues for detecting thermal stability changes of known drug targets upon *in vivo* dosing (*2*)(*10*), however, a strategy for the proteome-wide assessment of thermal stability and for target profiling *in vivo* has been lacking. Here we report a TPP strategy to assess thermal stability across tissue proteomes, henceforth referred to as tissue-TPP, and to study engagement of targets and off-targets in organs of animals dosed with the marketed HDAC inhibitor panobinostat.

To determine the thermal stability of tissue proteomes, three rats were sacrificed and biopsies derived from liver, lung, kidney, and spleen, were directly incubated for three minutes at a range of temperatures from 37° to 64° C. After mechanical tissue disruption with a bead-based procedure and addition of the mild detergent IPGAL and DNAse, the soluble protein fraction was analyzed by quantitative mass spectrometry resulting in a total of 11 032 proteins detected in the four organs (**fig. S1A, Data S1**). Proteome-wide thermal stability profiles recorded across the different organs showed very similar characteristics (Fig. 1A, **fig. S1B**) and were highly comparable to that generated with rat cell line, RBL-2H3 (Fig. 1B). This demonstrates that, despite the high diversity in structure and density of the different tissues, the heating process was homogeneous throughout the samples validating the procedure for proteome-wide thermal stability analysis in organs.

**Fig. 1.**
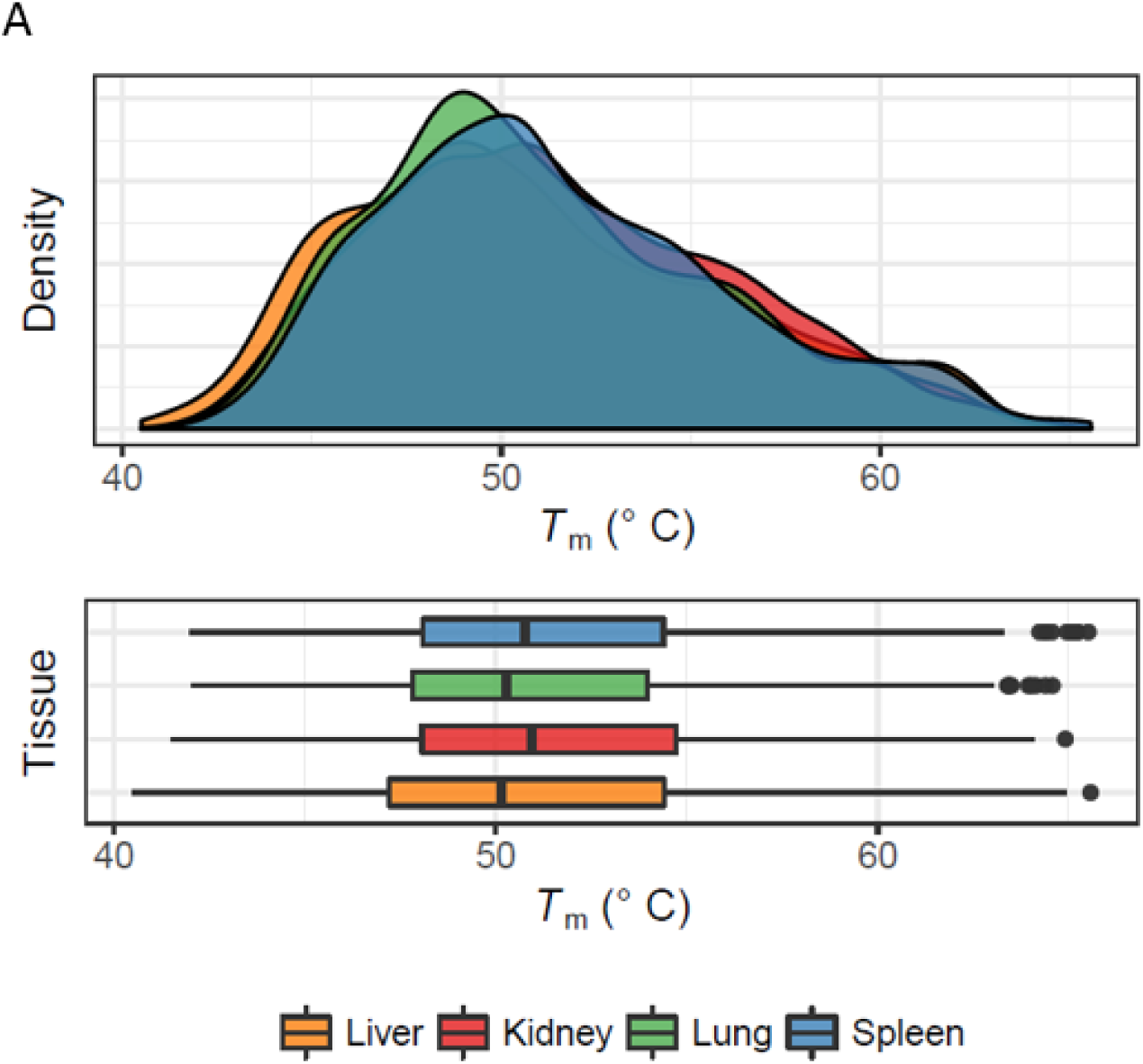
Analysis of proteome thermal stability in rat spleen, lung, kidney and liver biopsies. Freshly prepared tissue biopsies were exposed to temperatures spanning 37°C to 64°C for 3 min, disrupted using ceramic beads, and extracted. The soluble proteins were retrieved and analysed using quantitative multiplexed mass spectrometry. (A) Density plots and boxplots of protein melting temperatures (*T_m_*) determined by TPP in intact tissue biopsies. Plots represent the 1693 proteins for which melting points were determined in all four tissues. The median *T_m_* for each tissue is indicated in the boxplot by a black vertical line.

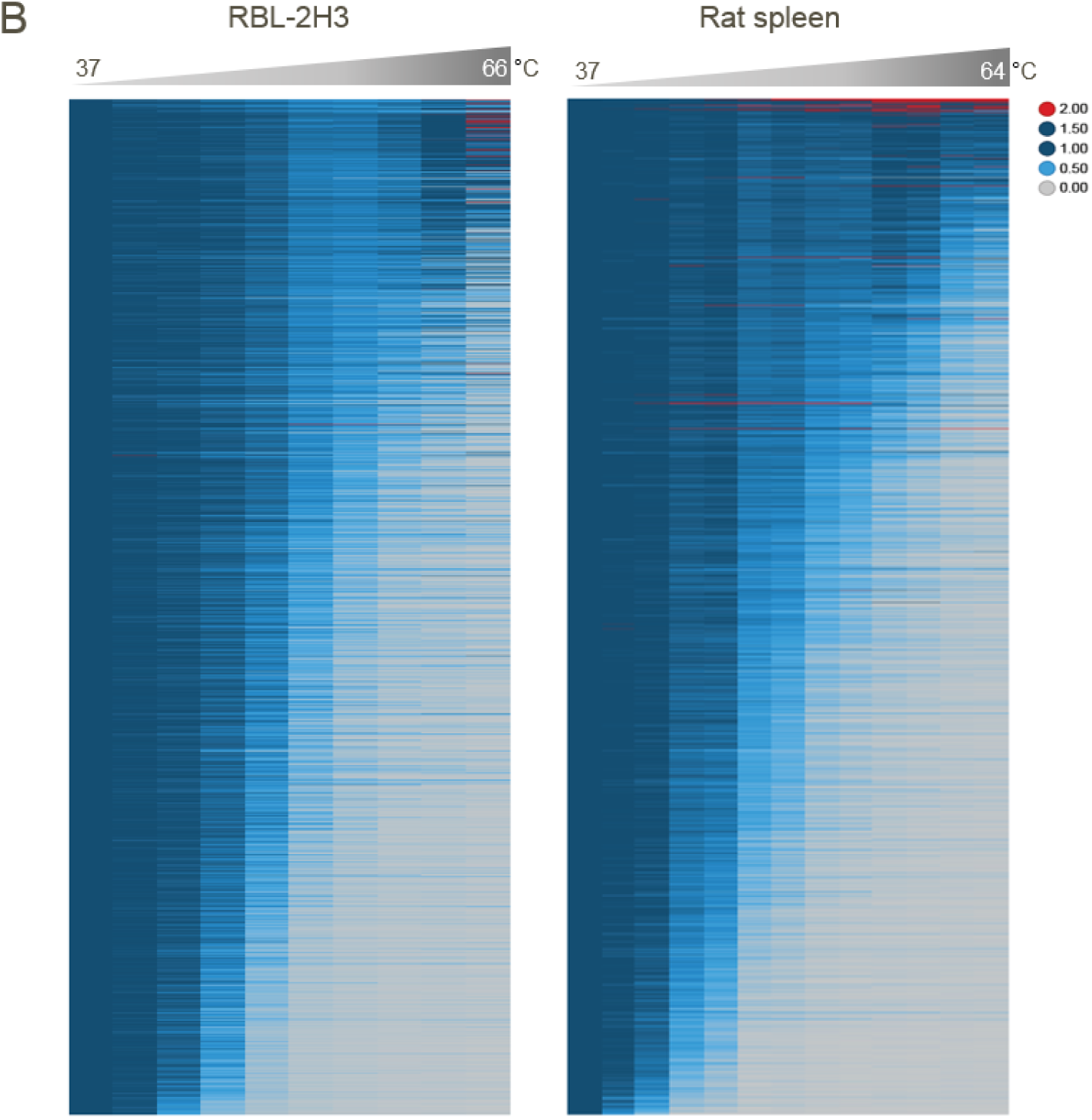
(B) Comparison of the thermal stability of soluble proteins in intact RBL-2H3 cells (left) and rat spleen biopsies (right), displayed as heatmap. The median relative abundance across all replicates (RBL-2H3 n=2; spleen biopsies n=6) at the indicated temperature is shown for each protein as fold-change relative to the lowest measured temperature (37°C). The 1739 proteins are plotted in the same order (See also Suppl. Fig. 1B).

Next, we performed pairwise comparisons of melting curves to detect proteins with different thermal stability across tissues using an F-test statistic-based strategy comparing the shapes of protein thermal denaturation curves (*11*) (**Data S2**). This made it possible to include incomplete denaturation curves in the analysis which were not interpretable with standard melting point based comparisons, thus increasing the number of analyzed proteins by 16-20% (**fig. S1C**, **Data S3**). A total of 637 proteins showed significantly different melting profiles between any of the four organs. Hierarchical clustering revealed greater similarities of thermal stability profiles in liver and kidney as well as between spleen and lung (Fig. 1C, **Data S4**). Several gene ontology (GO) terms were significantly enriched for proteins showing differential thermostability profiles (**fig. S1D**, **Data S5**). For example, five GO terms were related to fatty acid metabolism and proteins involved in fatty acid beta-oxidation, such as Acad9, Etfdh and Acadm, showed higher thermal stabilities in kidney and liver as compared to spleen and lung (Fig. 1C, **fig. S1E**, **table S1**). These differences could reflect the high capacity for fatty acid beta-oxidation of hepatocytes and proximal tubules in the kidneys (*12*). Other significantly enriched biological processes include amino acid activation and amino acid metabolism (**fig. S1D**, **table S2**) and for example proteins involved in the *cellular modified amino acid metabolic process*, such as Aldh7a1, Gatm and Gcdh were more thermostable in liver and kidney as compared to the other organs (Fig. 1D, 1E). Since the kidneys as well as the liver play a key role in the synthesis and interorgan exchange of several amino acids in the body (*13*), the increased thermal stability of enzymes involved in these processes could reflect different activity of the involved metabolic pathways. In order to validate this hypothesis, we quantified endogenous metabolites in these samples using an untargeted metabolomics approach (*14*). In total 5,031 ions were detected and these mapped to 2,363 metabolites (*15*) (**Data S6, S7, S8, fig. S1F**). Pathway enrichment analysis resulted in several pathways being significantly enriched between each pair of organs (**table S3**, **Data S9**). Amongst these, 12 amino acid metabolism related pathways showed profound differences between liver and kidney as compared to spleen and lung (**table S3**). Similar to thermal stabilities of proteins involved in the amino acid synthesis and degradation pathways, the metabolites associated with these pathways, such as the lysine degradation and arginine and proline metabolism pathways (Fig. 1F), were present at significantly lower levels in spleen and lung as compared to liver and kidney. In summary, in agreement with previous reports on cultured cells (*5*), trends observed in protein thermal stabilities across the tested organs were largely recapitulated by metabolite profiles due to metabolite based protein stabilization (*3*) and thus inform on the metabolic state in tissue.

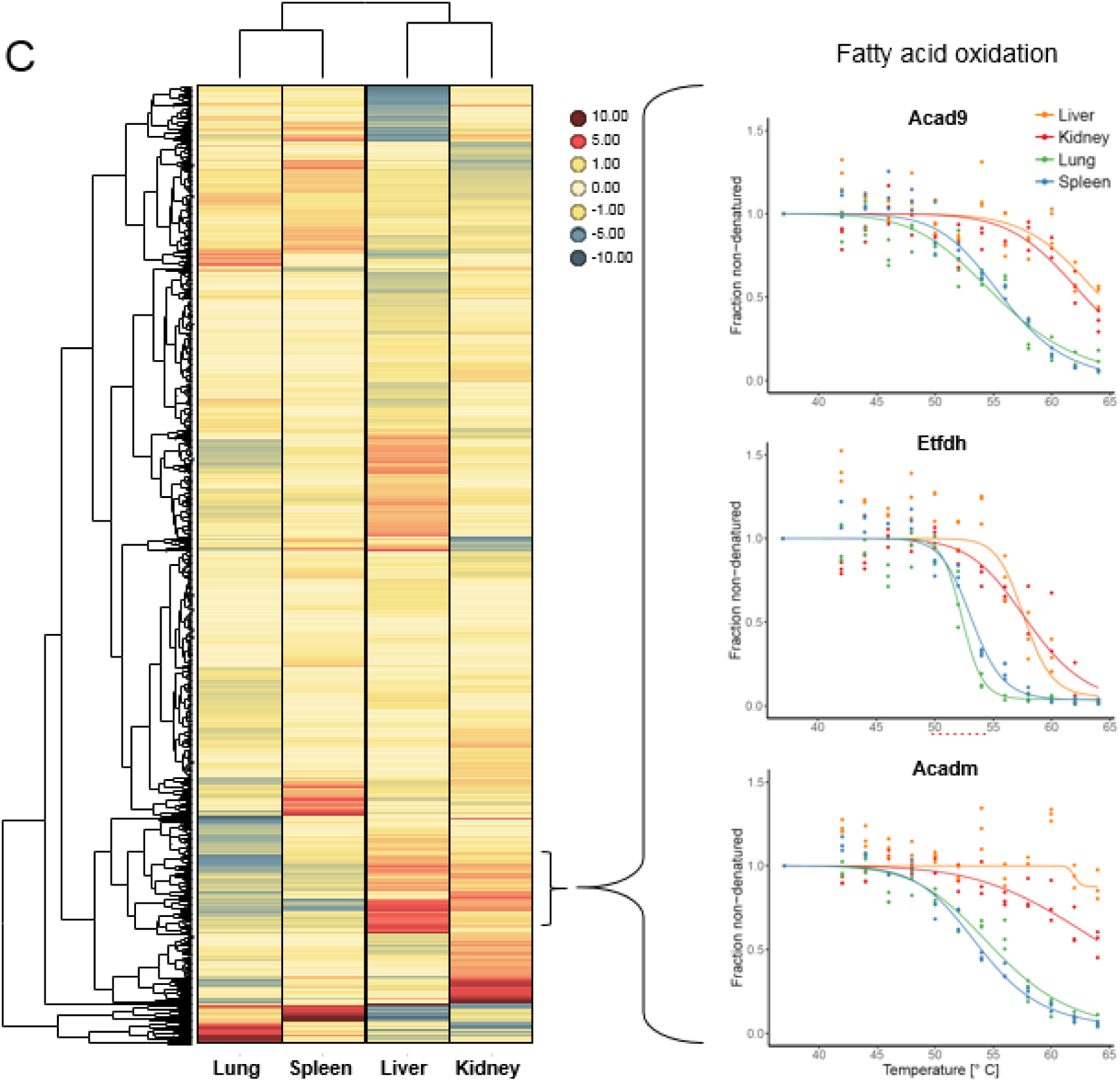
(C) Hierarchical clustering based on the median normalized AUCs of protein melting curves with significant differences between organs as detected in biopsies of vehicle-treated rats (637 proteins, left panel). Liver and kidney cluster together as well as lung and spleen. Proteins involved in fatty acid oxidation cluster together as indicated, and representative melting curves derived from three biological replicates are shown for three proteins (Acad9, Etfdh and Acadm) in four tissues (panels on the right). The fraction of denatured protein is normalised to the 37°C control and curves are fitted across the temperature range.

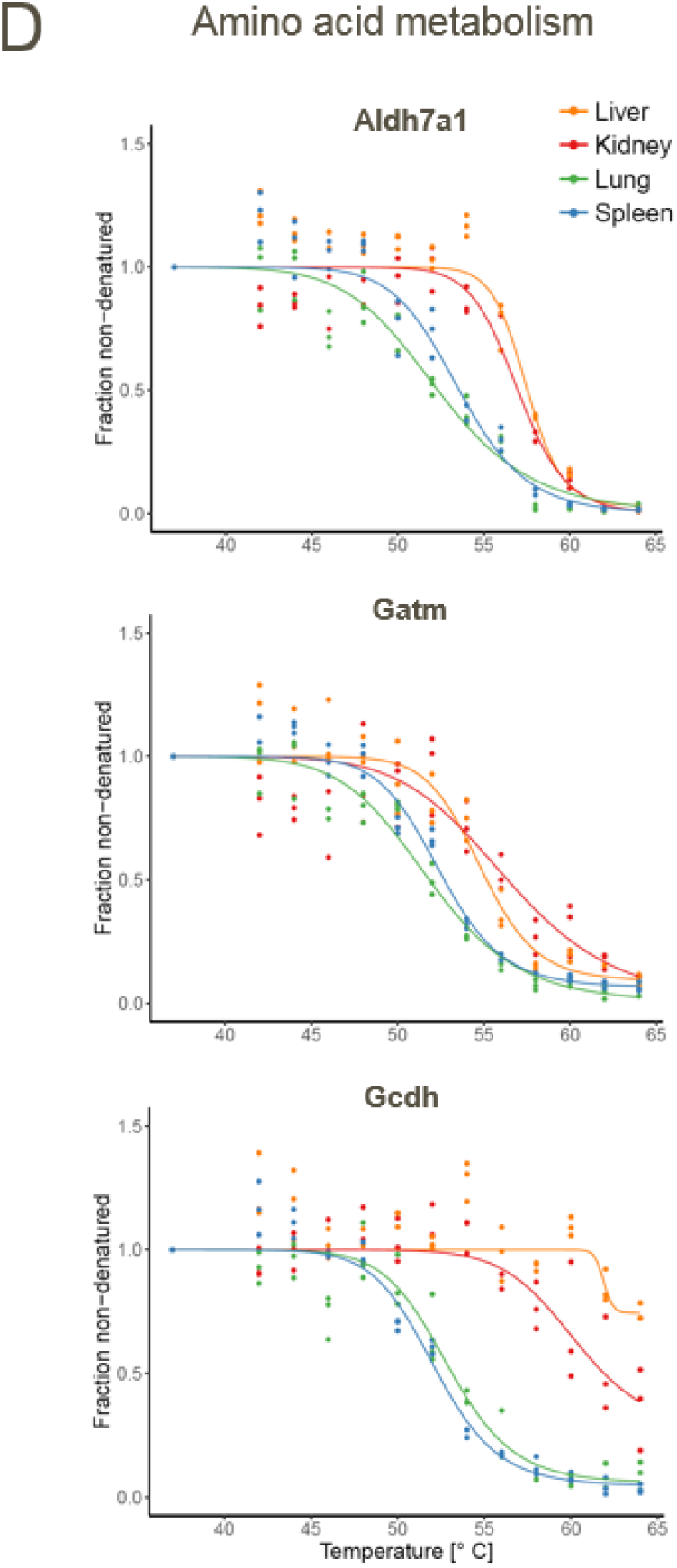
(D) Melting curves of three proteins involved in amino acid metabolism (Aldh7a1, Gatm and Gcdh) in tissue biopsies of vehicle-treated rats (as in panel 1C).

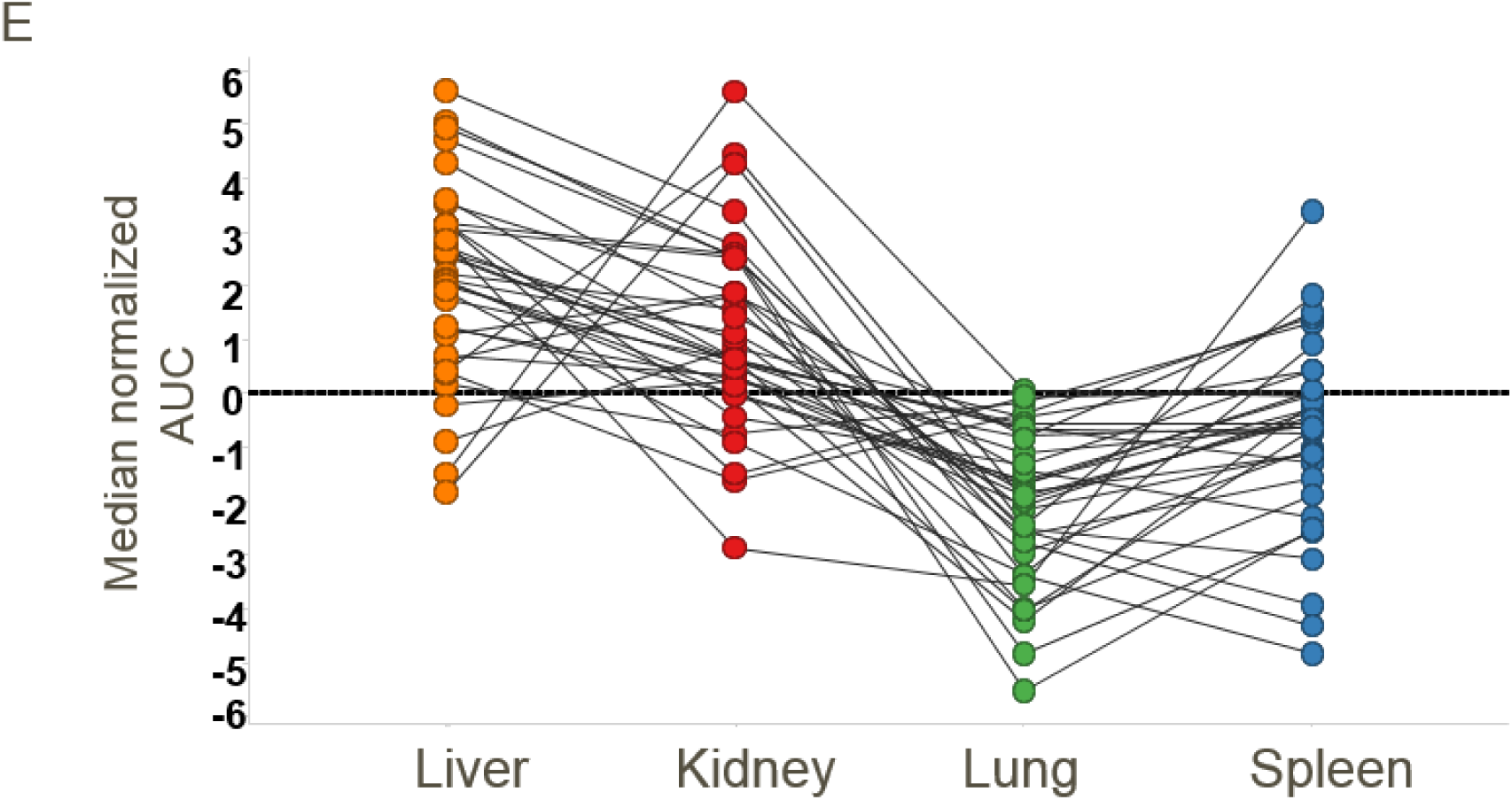
(E) Line chart of median normalized AUCs of melting curves for proteins involved in amino acid metabolism that are differentially stabilised in tissue biopsies of liver, kidney, lung or spleen from vehicle-treated rats.

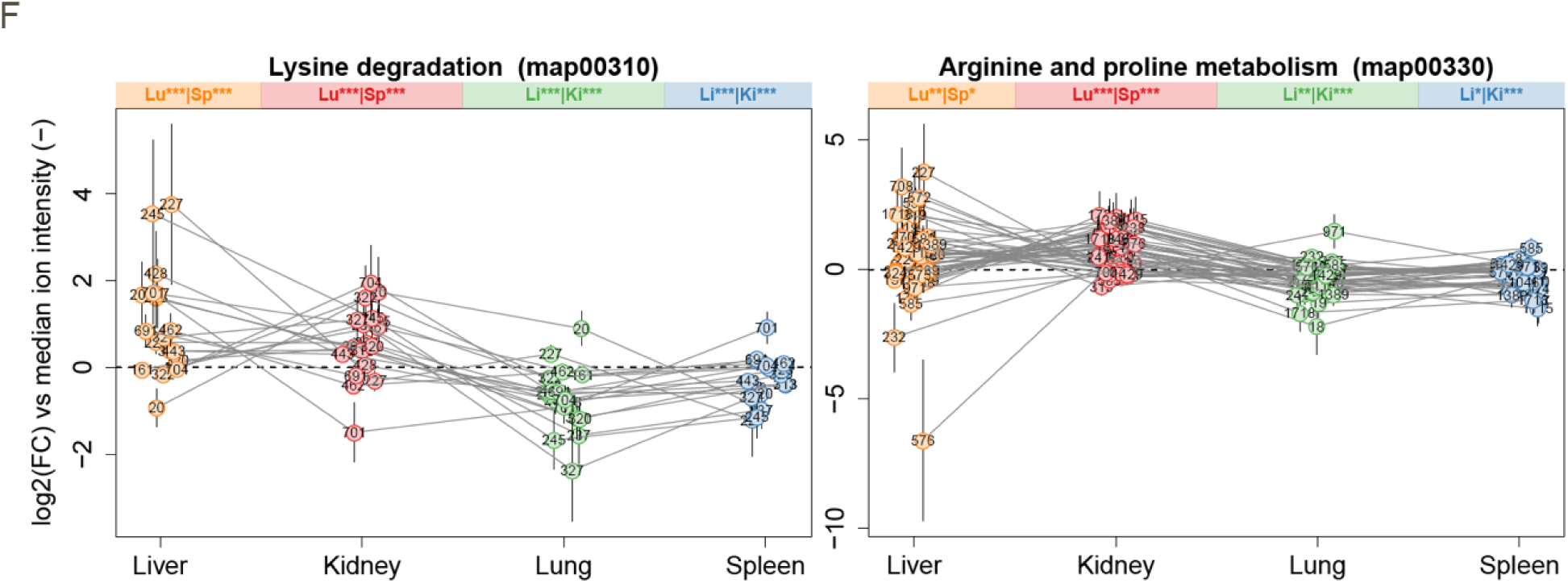
(F) Lysine degradation and arginine proline metabolism. Line charts show median-normalized intensities of metabolites across tissues in indicated metabolic pathways according to KEGG pathway definitions. Metabolites are identified by numbers as defined in **Data S8**. For each tissue, statistical significance of metabolite abundance distribution relative to other tissues is indicated above the plotting regions, *p<0.05, **p<0.01, ***p<0.001, otherwise p>0.05. P-values calculated using unpaired two-sided t-tests.

In addition to intracellular metabolite levels, several other factors can explain altered thermal stability of a protein, including post translational modifications and protein-protein interactions (*6*)(*7*). For example, we observed significant changes (p-value ≤ 0.001) in thermal stability for the signal transducers and activators of transcription (Stat) family of signaling proteins. These proteins were consistently less stable in liver compared to other organs (Fig. 2A). Since Stat signaling is also regulated by tyrosine phosphorylation, we hypothesized that the observed stability changes could reflect differences in phosphorylation state. Indeed, we found elevated levels of phosphorylated Stat5a/b in liver biopsies but not in kidney (**fig. S2A**), possibly reflecting the key roles of Stat proteins in the liver, for example for proliferation of the hepatocytes (*16*).

**Fig. 2.**
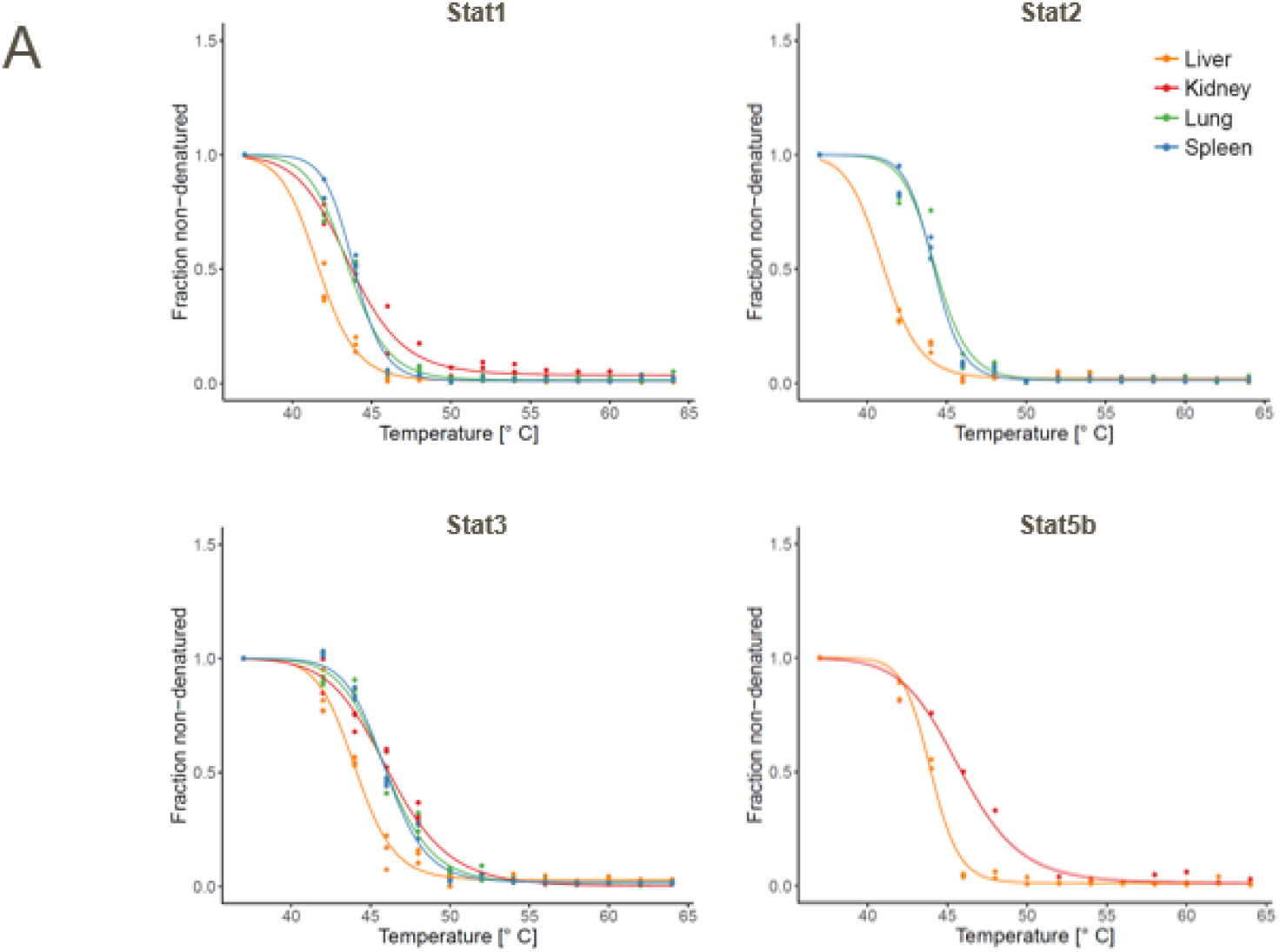
Comparison of the thermal stability of proteins associated with selected biological complexes or functions in rat spleen, lung, kidney and liver biopsies. (A) Melting curves of Stat1, Stat2, Stat3 and Stat5b in live kidney, liver, lung or spleen pieces.

Among the GO terms significantly enriched for cellular components, five were related to the proteasome (**fig. S2B**), indicating that also changes in supramolecular complexes could be detected in this tissue-TPP data set. We then analyzed the melting behavior of protein complexes in different organs using statistical methods previously reported for cultured and primary cells (*8*)(*7*)(*17*). We found that the melting points of complex subunits were significantly more similar than expected by chance and that several complexes showed different melting behaviors across the four tissues (**fig. S2C**, **Data S10 and S11**). Hierarchical clustering of 66 identified complexes recapitulated the organ clusters generated on a proteome level (Fig. 2B, 1C). We observed differential melting behavior of the 19S and the 20S proteasomal subunits in tissues, with the 20S core subunit having consistently higher thermal stability (Fig. 2C). Notably, the thermal denaturation of Psmd4, the only stoichiometric proteasomal subunit which was reported to be present in cells also outside this complex and that is required for the recognition of poly-ubiquitinylated proteins, was very different from the other subunits, in particular in lung tissue (*18*) likely reflecting different degrees of association with the 20S particle. Further, Psmd9, a chaperone involved in proteasome assembly, but not a constitutive member of the assembled proteasome complex, had systematically higher melting points in all tissues, suggesting a loose association to the 19S (*19*). Finally, Psmd5, a proteasomal member reported to act as proteasome inhibitor (*20*), showed the lowest thermal stability in liver, lung, and spleen, while it co-melted with the other 19S components in kidney, suggesting an organ specific regulation of the 19S complex in kidney (Fig. 2C).

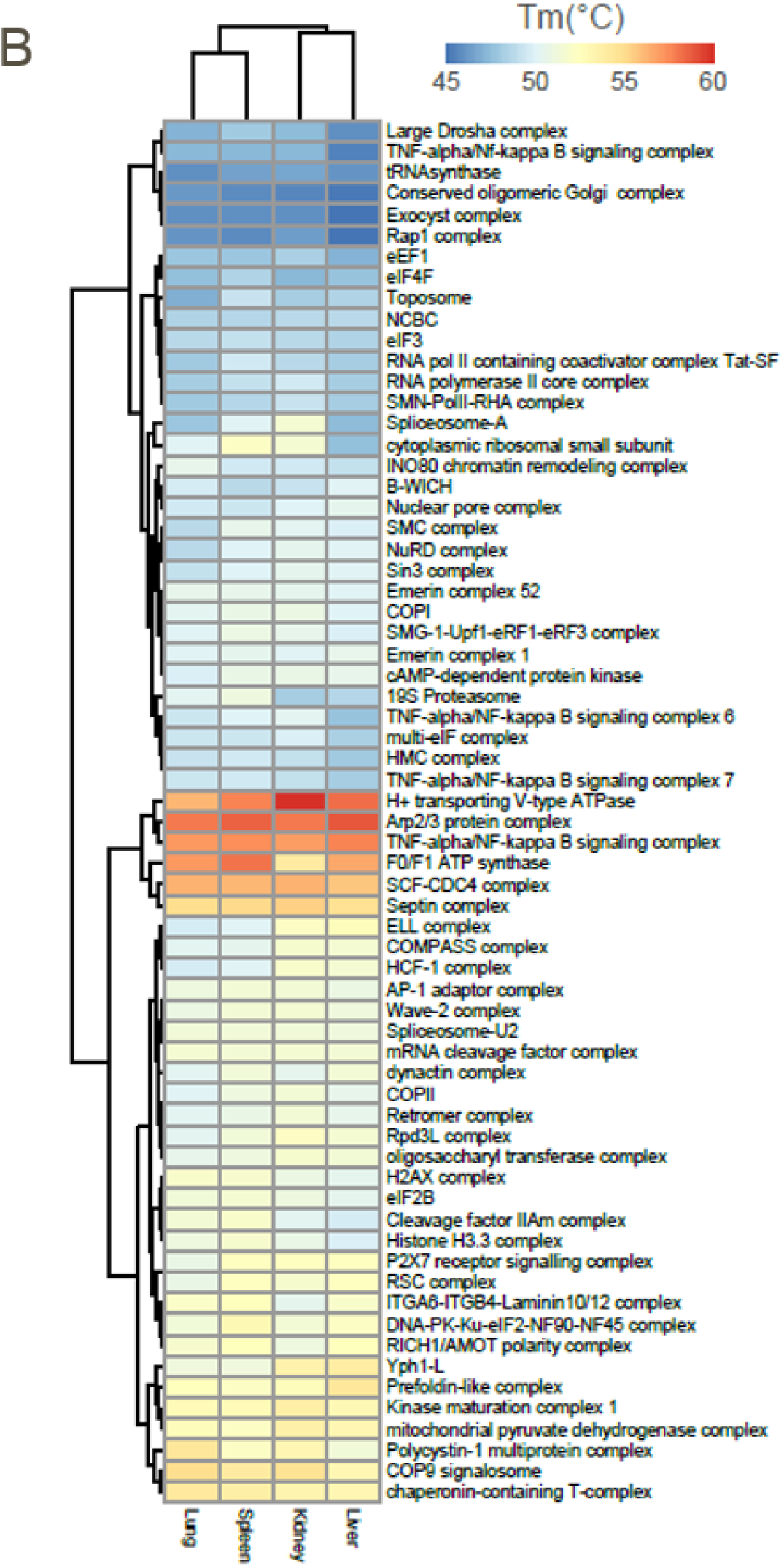
(B) Heatmap and hierarchical clustering analysis of protein complexes based on the median melting points of subunits identified across tissues.

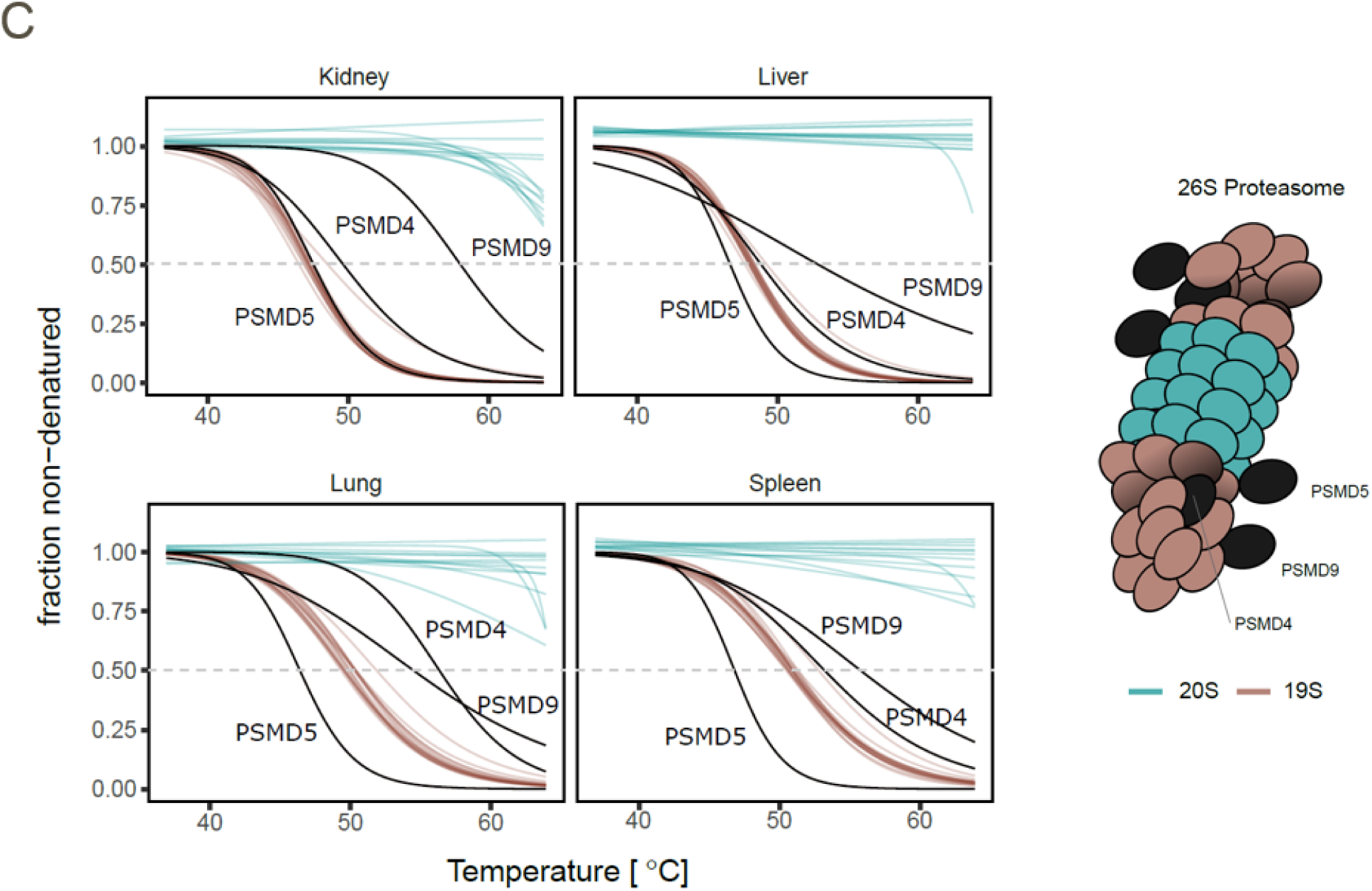
(C) Simplified 26S proteasome structure (right panel) and melting curves (left panel) for identified proteasome 19S and 20S complex members.

The H+ transporting V-type ATPase, known to be required in kidney for the deacidification of the blood (*21*), had a significantly higher median melting point specifically in this organ (**Data S10**). Detailed comparison of the individual complex subunits revealed that along with an overall increase in thermal stability of the V1 subcomplex members, the melting point of the C subunit of V1 (v1c1) was significantly increased in kidney compared to the other tissues (Fig. 2D). This is indicative of a higher population of fully assembled complexes in kidney as the C subunit is known to dissociate from V1 upon detachment of V1 from V0, providing a rationale for the higher thermal stability measured for the complex in this organ (Fig. 2D) (*22*).

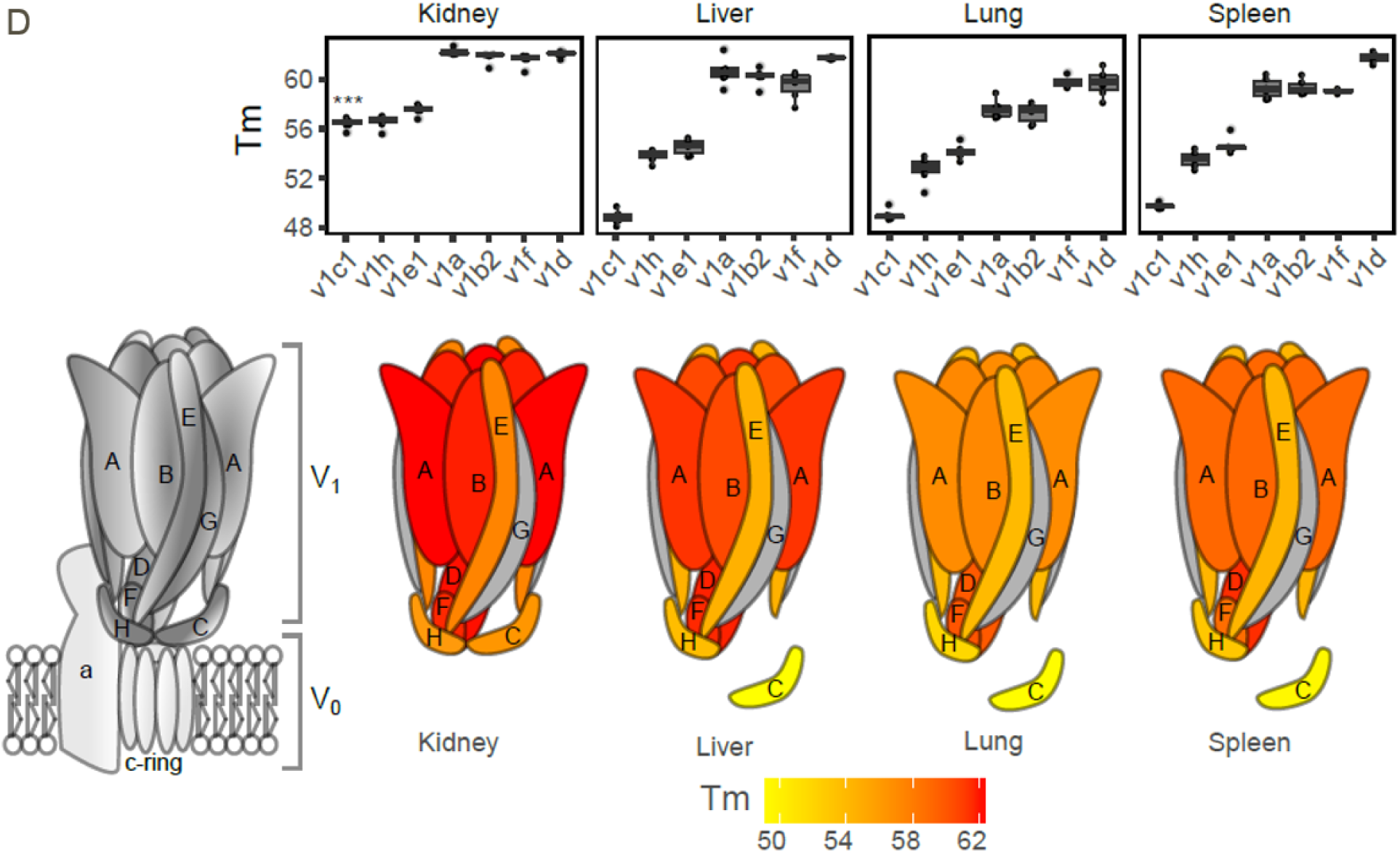
(D) Boxplots of melting points determinedfor the different subunits of the V1 subcomplex of V-type ATPase across tissues (top panel) and melting points of the V-type ATPase mapped onto its simplified structure (lower panel).

Thus, differences in protein thermal stability measured by tissue-TPP reflect the known structural and functional differences of protein subunits of complexes.

Organ specific features were also observed for the ribosome. The cytoplasmic small ribosomal subunit showed a lower stability in liver as compared to the other organs (**fig. S2D**). This finding could be explained with the previously reported circadian regulation of ribosome biogenesis in the liver (*23*), and the recent report that thermal stabilities of the ribosomal subunits are lower when they are in an unassembled state, e.g. during synthesis (*8*). Similarly, the eIF2B translation initiation complex and the eEF1 translation elongation complex showed lower T_m_ specifically in liver, possibly reflecting the cyclic regulation of protein translation taking place in this organ (*24*) (**fig. S2E**, **S2F**).

Also, the thermal denaturation of the nuclear pore complex member Ranbp2 differed from the other subunits particularly in liver but not in kidney (**fig. S2G**). This observation might reflect different activities of Ranbp2 in the analyzed organs, e.g. mediated by the binding of this subunit to other proteins as previously observed in hepatocytes (*25*). Finally, reduced thermal stability of the COP9 signalosome (CSN) observed in liver might be linked to the reported role of this complex in the proliferation of hepatocytes (*26*) (**fig. S2H**).

In summary our data demonstrate that tissue-TPP recapitulates physiological processes as well as differences related to energy metabolism, signaling or protein homeostasis in organs.

Next, we devised a tissue-TPP strategy for proteome-wide measurements of drug target engagement upon ex-vivo dosing of tissue samples and directly in dosed animals. The *in vitro* to *in vivo* translation of the molecular mode-of-action of a drug is often extremely challenging and the ability to measure target and off-target engagement *in vivo* is critical for understanding efficacy but also adverse reactions already in early preclinical phases (*26*). Here, we investigated drug-protein interactions of the marketed HDAC inhibitor panobinostat in organs thereby taking advantage of the extensive *in vitro* characterization of its selectivity in biochemical and cell-based assays (*27*)(*5*). First, we compared protein target binding of panobinostat in cell lines with tissues using a crude extract protocol (*4*). In this lysis strategy, cells are broken up mechanically in the absence of detergent. Protein extracts containing cellular membranes and other insoluble structures are then heated to different temperatures and subsequently extracted with a mild detergent. When comparing membrane proteins detected in HepG2 crude extracts and intact cells, we observed similar amounts but detected approximately four-fold less in detergent-free extract (**fig. S3A**, **Data S12**). Further, the distribution of melting temperatures between intact cells and crude extracts was comparable (**fig. S3B**) and similar results were also observed in crude extracts from three additional rat organs, kidney, lung and spleen (**fig. S3C**, **Data S13**). We next performed two-dimensional (2D-) TPP (*3*) with panobinostat in crude extracts derived from the HepG2 cell line, mouse and rat liver tissue and found highly comparable target profiles across all extracts (**fig. S3D, S3E, Data S14, table S4**). Notably, the multi-pass membrane, endoplasmic reticulum localized protein FADS1, only identified and stabilized in cells and not detected in PBS extracts TPP (*5*) (**fig. S3F**), was stabilized in crude extract, thus indicating that this protein is a direct target of panobinostat. This data shows that crude extracts derived from animal tissues can be used to generate *in vitro* cross-species selectivity profiles of small molecules.

Next, we adopted the tissue-TPP strategy to biopsies freshly prepared from sacrificed animals treated with compound *in vitro*. Biopsy punches of 4 mm diameter from rat spleen, were incubated with panobinostat for 2.5 h, heated at a temperature range from 42 °C to 63.9 °C and analyzed as described (Fig. 3A). Significant stabilization of known panobinostat targets such as Hdac1, Hdac2, Hdac10 and Tt38 was observed. In addition, Mier1, a known Hdac1/2 interactor(*27*) was also stabilized, suggesting an indirect effect mediated by the stabilization of the Hdacs 1 and 2. Interestingly, the melting profile of the dehydrogenase/reductase SDR family member 1, Dhrs1, was significantly altered following treatment with panobinostat (Fig. 3B, **fig. S3G**, **Data S15**, **S16**). As this experiment was performed in live tissue, the effect could result either from the direct interaction of a drug or a drug metabolite with the protein or caused by an indirect effect downstream of panobinostat targets.

**Fig. 3.**
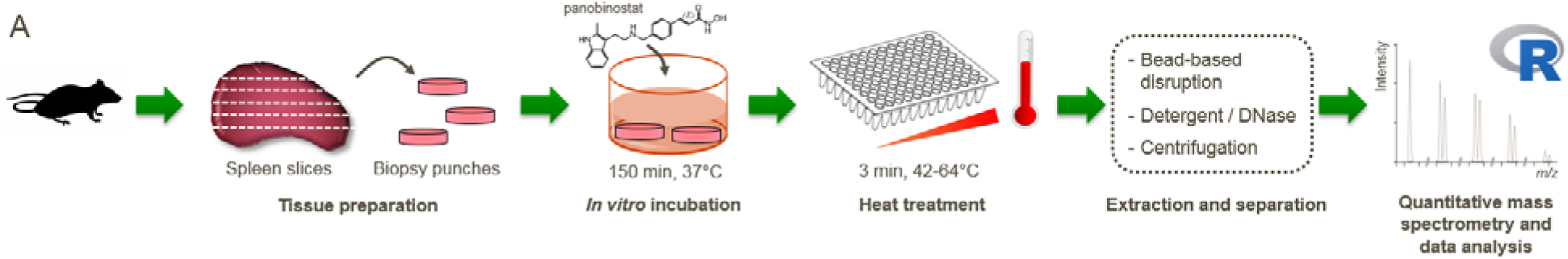
Target identification and occupancy measurements of the HDAC inhibitor panobinostat in tissues and blood. (A) *In vitro* “tissue-TPP” workflow for rat spleen. Biopsy punches (4 mm, 2 per well) generated from fresh spleen precision cut slices (thickness 500 μm) of untreated rats were treated with panobinostat for 150 min in cell culture media at 37°C, 5%CO_2_, and then transferred into PCR plates. After heat treatment over 12 temperatures for 3 min in buffer and one freeze-thaw cycle, the tissues were disrupted using ceramic beads and incubated with detergent and DNase for 60 min at 4°C. The soluble proteins were separated by centrifugation and identified and quantified using mass spectrometry.

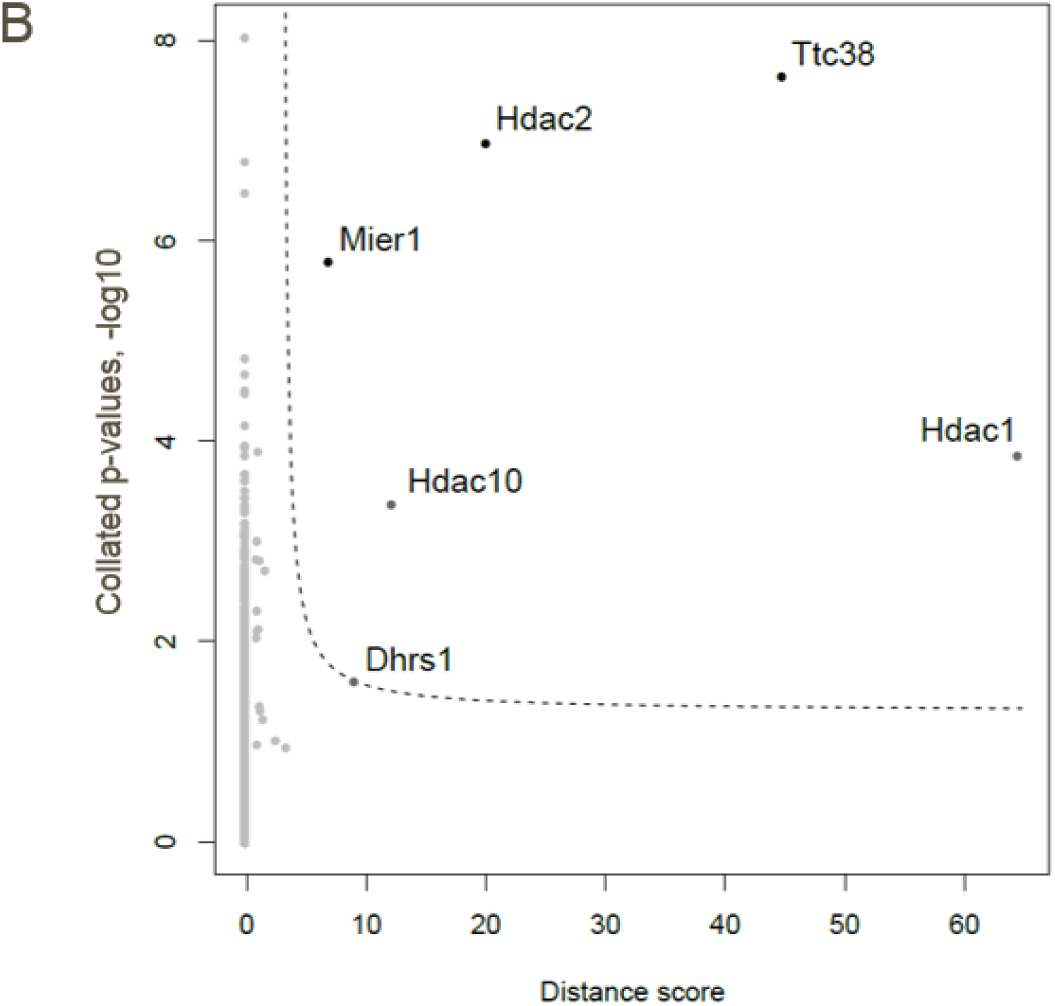
(B) Volcano plot of protein stability changes and corresponding p-values measured by TPP in rat spleen biopsies upon in vitro incubation with panobinostat (as described in A). p-values were calculated against the vehicle condition for each protein, temperature and concentration, and then collated. A distance score was calculated by collating protein abundance ratio of the panobinostat/vehicle conditions, thus reflecting the aggregated change in protein stability. Black dots: proteins significantly changed after Benjamin-Hochberg (BH) correction, dark grey dots: proteins not significantly changed after BH correction, light grey dots: proteins not significantly changed.

Finally, we then applied tissue-TPP to organ biopsies derived from rats administered intravenously with 10 mg/kg panobinostat and sacrificed 90 minutes post-dose. After preparation of tissue biopsies from kidney, liver, lung and spleen, these were incubated at a range of temperatures and then analyzed by multiplexed quantitative mass spectrometry (Fig. 3C, **fig. S3H**, **Data S1**). Significant stabilization of Hdac1, Hdac2 and Tt38 was observed in all analyzed tissues (Fig. 3D, **Data S17**). Hdac6 was stabilized in both lung and spleen, while stabilization of Fads1, could be observed exclusively in lung. Some panobinostat off-targets identified in the crude extracts (**fig. S3D**), were either not detected in tissue samples because of low expression or stabilization was not significant at the drug exposure achieved in these tissues. Interestingly, while Hdac1, Hdac2 and Ttc38 were less stabilized in liver where the lowest free panobinostat concentrations were observed, the effect on Dhrs1 was highest in this organ as compared to the other organs (Fig. 3E, **table S5**). Lower Hdac engagement in the liver is in agreement with previous reports on rapid metabolization of panobinostat (*28*). We further hypothesized that the effect observed on Dhrs1 could be a result of drug metabolization rather than direct binding to panobinostat and since it was also observed in *in vitro* treated spleen biopsies, we investigated whether panobinostat is metabolized in these samples. The detection of the amide metabolite M37.8 formation (*5*) (**fig. S3I**) suggests that Dhrs1 stabilization could be related to panobinostat metabolization.

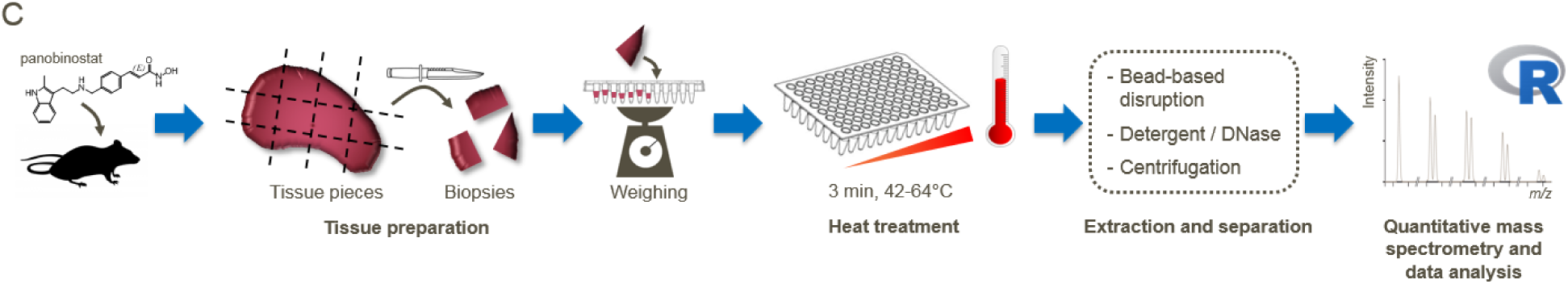
(C) “Tissue-TPP” workflow for tissue biopsies from *in vivo* dosed rats. Rats were dosed intravenously with panobinostat or vehicle. 1.5 hr post-injection, the animals were sacrificed, and lungs, spleen, liver and kidney were dissected into pieces Biopsies were weighed and transferred into PCR plates. After heat treatment over 13 temperatures from 37 to 64°C for 3 min in buffer and one freeze-thaw cycle, the tissues were disrupted using ceramic beads and incubated with detergent and DNase for 60 min at 4°C. The soluble proteins were separated by centrifugation and identified and analyzed by quantitative mass spectrometry.

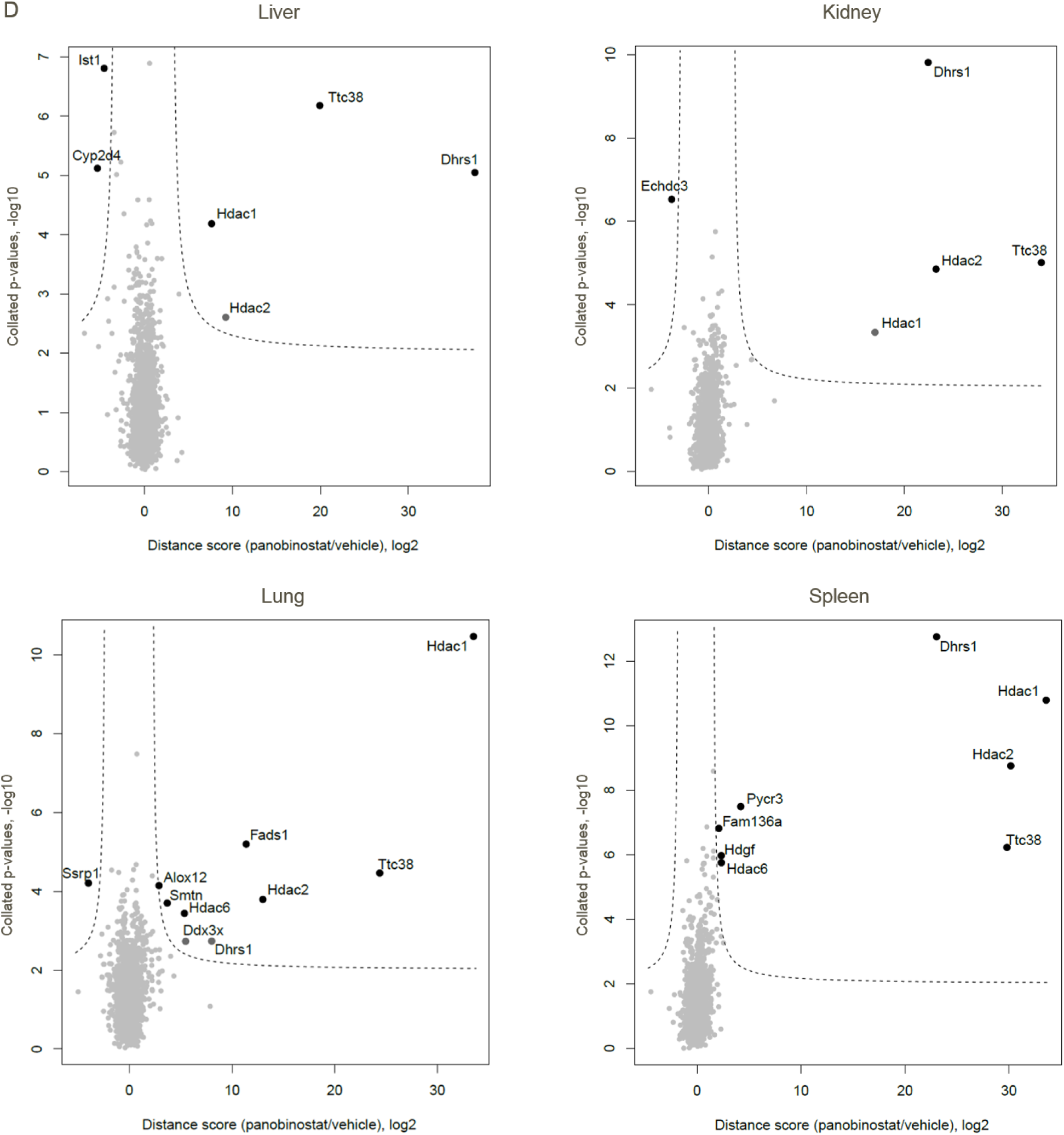
(D) Volcano plots with collated p-values (log10) and distance scores (collated log2 ratios) of proteins quantified in panobinostat-treated animals (n=3) vs vehicle treated animals (n=3) in liver, kidney, lung and spleen. Black dots: proteins significantly affected after Benjamin-Hochberg (BH) correction, dark grey dots: proteins not significantly affected after BH correction, light grey dots: proteins not significantly affected.

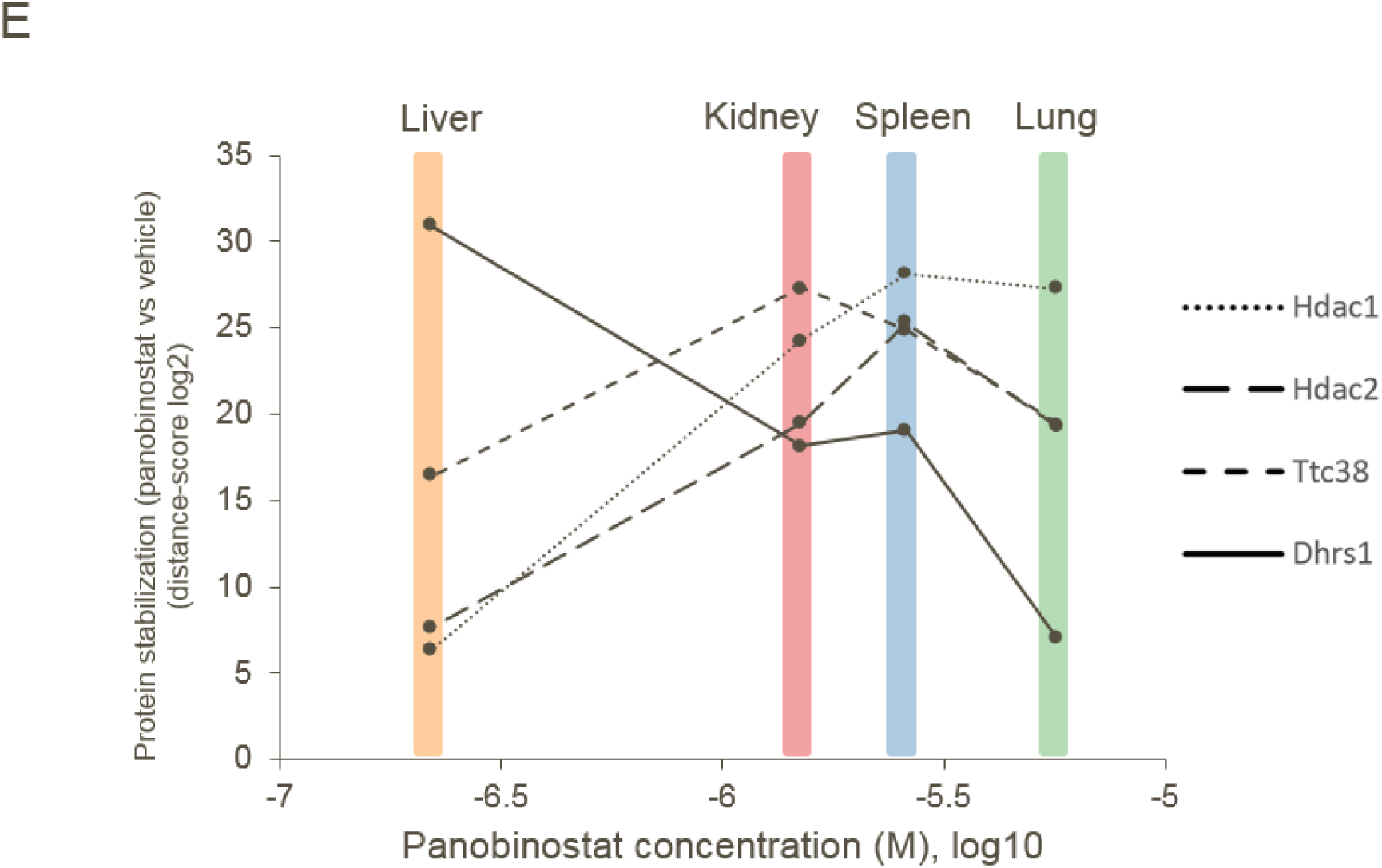
(E) Line charts plotting the distance scores of melting curves for in vivo panobinostat dosed vs. vehicle treated animals for Hdac1, Hdac2, Ttc38 and Dhrs1 against the total panobinostat concentrations measured in the respective tissues.

Overall, this data indicates that tissue-TPP enables the identification of targets and off-targets in tissue derived from animals dosed with non-covalent small molecule inhibitors and that the engagement measured for these targets correlates with drug exposure in tissue.

Monitoring target engagement in clinical studies enables optimizing dosing regimen to maximize efficacy while avoiding adverse effects. To enable target engagement measurements in samples readily accessible in clinical studies, we developed a thermal shift assay in whole blood, blood-CETSA. Freshly derived blood samples from rat, were treated with panobinostat or vehicle, aliquoted and subjected to heat treatment. After red blood cells removal, cells were lysed and protein aggregates were removed before protein detection was performed by antibody (Fig. 3F). In this set up, panobinostat induced dose-dependent stability changes of Hdac2, with a pEC50 of 7 (Fig. 3G). In experiments with human whole blood, a pEC_50_ of 7.1 was measured for Hdac2 (Fig. 3H, **fig. S3J**). Two additional marketed Hdac inhibitors, belinostat and romidepsin, were tested in human blood-CETSA and the measured pEC_50_s reflected the compound potencies for inhibiting Hdacs (*29*), further validating the assay (Fig. 3H).

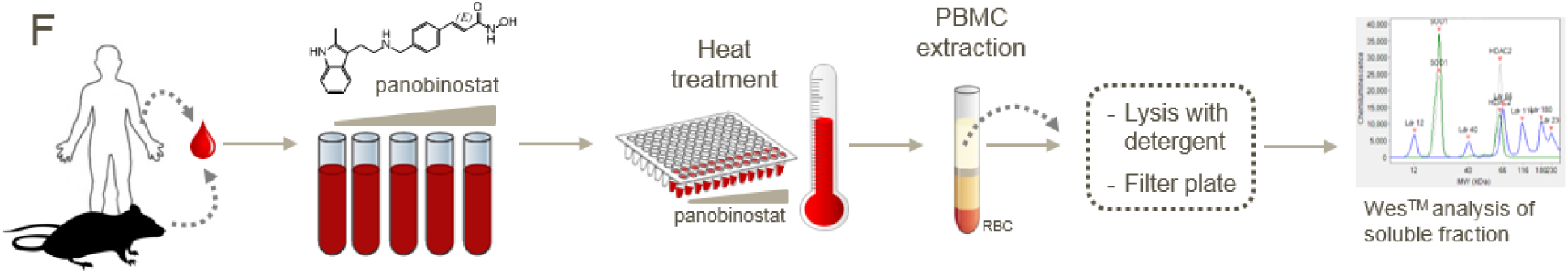
(F) Blood CETSA workflow. Fresh human or rat whole blood was treated *ex vivo* with panobinostat (90 min, 37°C) and directly heated in PCR plates (3 min) with subsequent separation of red blood cells (RBC). After lysis and filtering, direct target engagement in blood was measured by WES™ analysis of the soluble fraction.

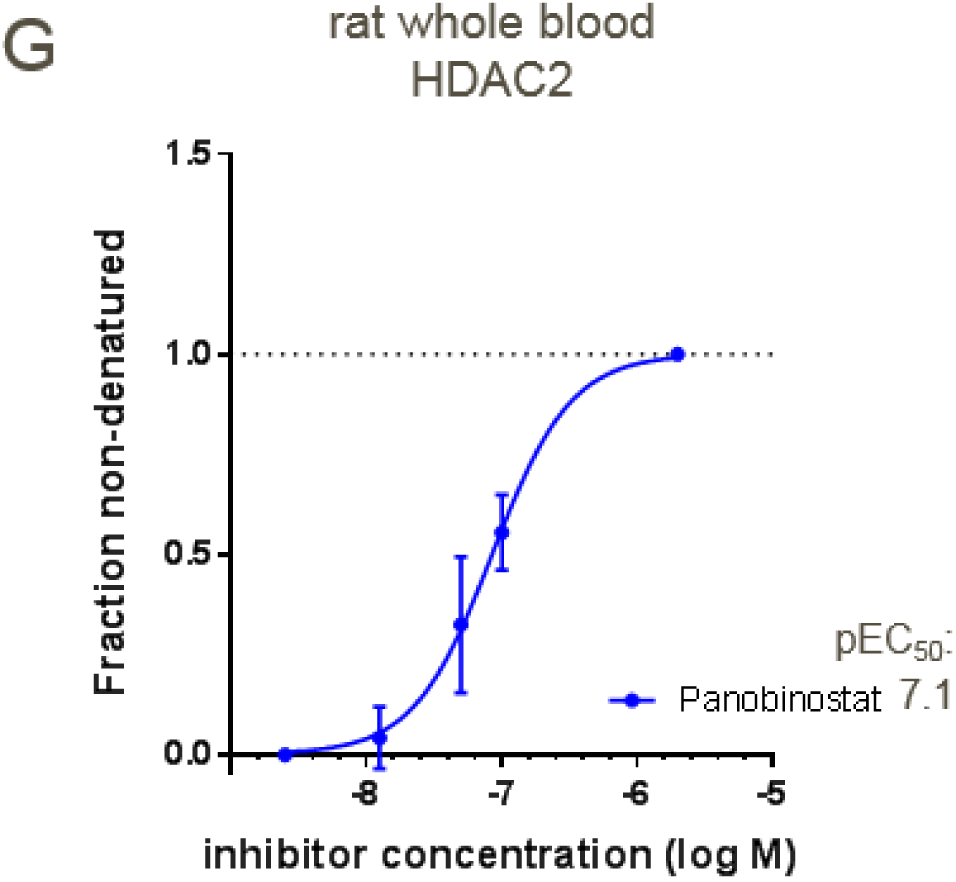
(G) Dose dependent stabilization of HDAC2 by panobinostat in rat blood measured with blood CETSA (n=4 rats, SEM shown). The fraction of denatured protein is normalised to the vehicle treated control.

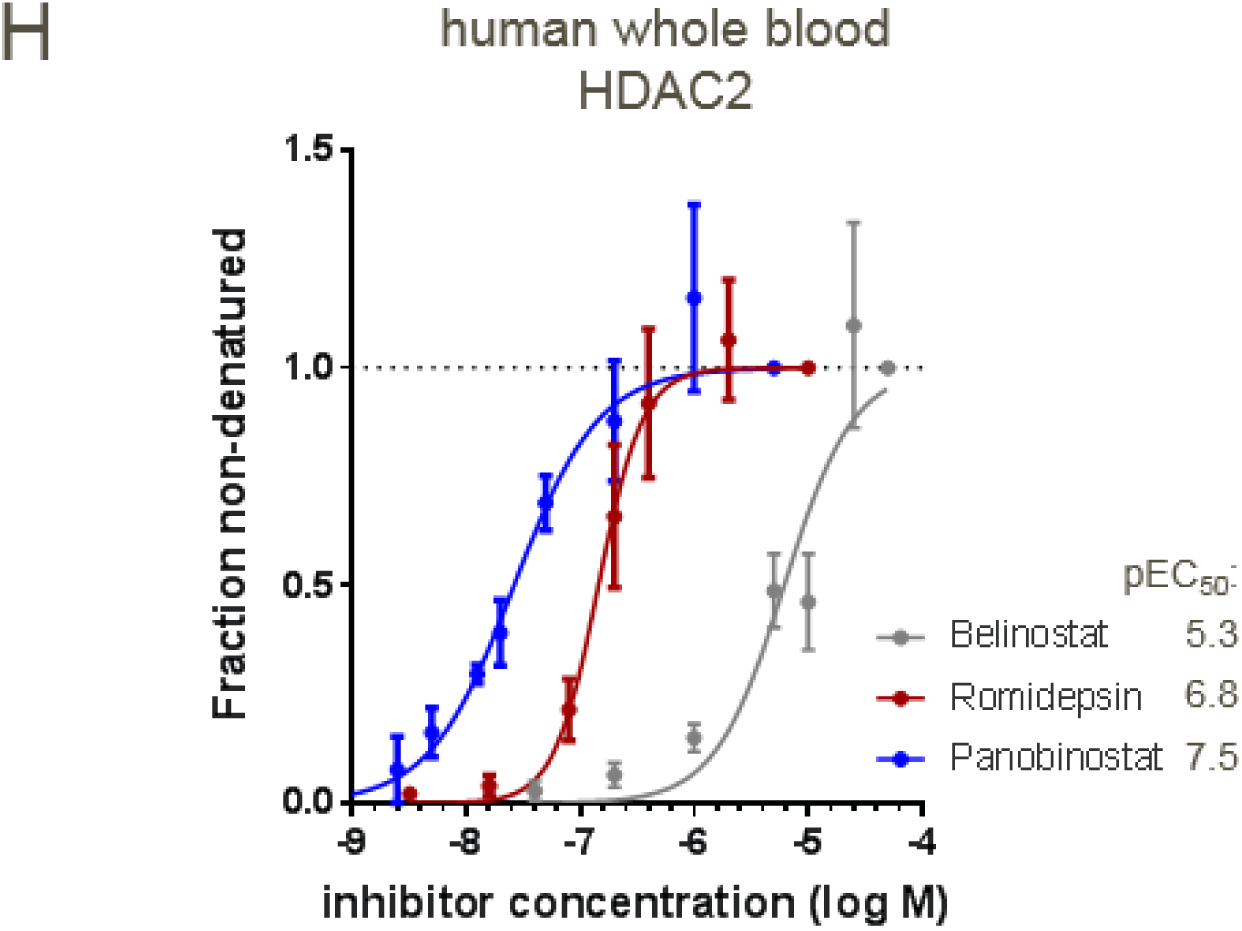
(H) Comparison of dose -dependent HDAC2 stabilization by the HDAC inhibitors panobinostat, belinostat and romidepsin for stabilisation of HDAC2 in human blood measured with blood CETSA (n=3 donors in duplicates, SEM shown), enabling ranking of compounds according to their affinities. The fraction of denatured protein is normalised to the vehicle treated control and curves are fitted as described in methods.

In clinical and pre-clinical studies, blood-CETSA could provide a quantitative measurement of the levels of target engaged after *in vivo* administration by the aid of compound spiked-in controls which would enable to normalize each individual blood sample (**fig. S3K**).

Our study provides a comprehensive map of proteome thermal stability in different organs. Differences in protein thermal stability between tissues inform on metabolic and signaling states, as well as complex assembly, thus providing a valuable resource to investigate physiological processes in the body. We also show that tissue-TPP can discover drug (off-)targets in an unbiased way following *in vivo* drug treatment and could inform on downstream effects, including consequences of drug metabolization. Finally, the blood-CETSA approach, could readily monitor drug target engagement in blood samples from *in vivo* and clinical studies. We foresee broad future use of these methods both in fundamental research as well as in preclinical and clinical settings.

## Acknowledgments

The authors thank Jürgen Stuhlfauth, Nilma Garcia-Altrieth, Kerstin Beß, Letizia Vitali and Bianca Dlugosch for the support in cell culture and lysates production; Markus Bösche, Tatjana Rudi, Manuela Klös-Hudak, Kerstin Kammerer and Michael Steidel for assistance with mass spectrometry; Stephan Gade for support in data uploading; Christopher Schofield, Anita Naidoo, Tracy Walker, Cerys Lovatt, Gerard Drewes, Paola Grandi and Friedrich Reinhard for scientific discussions and support; Edwige Nicodeme and Florence Blandel (Oncodesign, Les Ulis, France) for the excellent technical support for the animal study.

## Author contributions

JP, TW, DS, JV, CR, JP, ES, KS, JK, BH and DP performed experiments and analyzed data for Tissue-TPP; AR, KS and BH performed the experiments and analyzed the data for the Blood-CETSA; NK, DDC, MMK, MFS and DCS performed the statistical analysis; DT, WH and CHE gave scientific and experimental advice; GB, MB, JP, TW, AR and MMS designed experiments and analyzed data; GB, MB and MMS wrote the manuscript; GB and MB conceptualized the work.

## References

1. M. Bantscheff, S. Lemeer, M. M. Savitski, B. Kuster, Quantitative mass spectrometry in proteomics: critical review update from 2007 to the present. Anal. Bioanal. Chem. 404, 939–965 (2012).

2. D. M. Molina et al., Monitoring Drug Target Engagement in Cells and Tissues Using the Cellular Thermal Shift Assay. Science. 341, 84–87 (2013).

3. M. M. Savitski et al., Tracking cancer drugs in living cells by thermal profiling of the proteome. Science. 346, 1255784 (2014).

4. M. M. Savitski et al., Multiplexed Proteome Dynamics Profiling Reveals Mechanisms Controlling Protein Homeostasis. Cell. 173, 260–274.e25 (2018).

5. I. Becher et al., Thermal profiling reveals phenylalanine hydroxylase as an off-target of panobinostat. Nat. Chem. Biol. 12, 908–910 (2016).

6. F. B. M. Reinhard et al., Thermal proteome profiling monitors ligand interactions with cellular membrane proteins. Nat. Methods. 12, 1129–1131 (2015).

7. C. S. H. Tan et al., Thermal proximity coaggregation for system-wide profiling of protein complex dynamics in cells. Science. 359, 1170–1177 (2018).

8. I. Becher et al., Pervasive Protein Thermal Stability Variation during the Cell Cycle. Cell (2018), doi:10.1016/j.cell.2018.03.053.

9. L. Dai et al., Modulation of Protein-Interaction States through the Cell Cycle. Cell (2018), doi:10.1016/j.cell.2018.03.065.

10. T. Ishii et al., CETSA quantitatively verifies in vivo target engagement of novel RIPK1 inhibitors in various biospecimens. Sci. Rep. 7, 13000 (2017).

11. D. Childs et al., Non-parametric analysis of thermal proteome profiles reveals novel drug-binding proteins. submitted.

12. P. Bhargava, R. G. Schnellmann, Mitochondrial energetics in the kidney. Nat. Rev. Nephrol. 13, 629–646 (2017).

13. M. C. G. van de Poll, P. B. Soeters, N. E. P. Deutz, K. C. H. Fearon, C. H. C. Dejong, Renal metabolism of amino acids: its role in interorgan amino acid exchange. Am. J. Clin. Nutr. 79, 185–197 (2004).

14. T. Fuhrer, D. Heer, B. Begemann, N. Zamboni, High-throughput, accurate mass metabolome profiling of cellular extracts by flow injection-time-of-flight mass spectrometry. Anal. Chem. 83, 7074–7080 (2011).

15. D. S. Wishart et al., HMDB 4.0: the human metabolome database for 2018. Nucleic Acids Res. 46, D608–D617 (2018).

16. B. Gao, H. Wang, F. Lafdil, D. Feng, STAT proteins – key regulators of anti-viral responses, inflammation, and tumorigenesis in the liver. J. Hepatol. 57, 430–441 (2012).

17. T. Mathieson et al., Systematic analysis of protein turnover in primary cells. Nat. Commun. 9, 689 (2018).

18. R. Piterman et al., VWA domain of S5a restricts the ability to bind ubiquitin and Ubl to the 26S proteasome. Mol. Biol. Cell. 25, 3988–3998 (2014).

19. T. Kaneko et al., Assembly pathway of the Mammalian proteasome base subcomplex is mediated by multiple specific chaperones. Cell. 137, 914–925 (2009).

20. S. M. Shim et al., Role of S5b/PSMD5 in proteasome inhibition caused by TNF-α/NFĸB in higher eukaryotes. Cell Rep. 2, 603–615 (2012).

21. Q. Al-Awqati, Plasticity in epithelial polarity of renal intercalated cells: targeting of the H(+)-ATPase and band 3. Aw. J. Physiol. 270, C1571–1580 (1996).

22. P. M. Kane, Disassembly and reassembly of the yeast vacuolar H(+)-ATPase in vivo. J. Biol. Chem. 270, 17025–17032 (1995).

23. C. Jouffe et al., The circadian clock coordinates ribosome biogenesis. PLoSBiol. 11, e1001455 (2013).

24. F. Atger et al., Circadian and feeding rhythms differentially affect rhythmic mRNA transcription and translation in mouse liver. Proc. Natl. Acad. Sci. U. S. A. 112, E6579–6588 (2015).

25. D.-H. Kim et al., Critical role of RanBP2-mediated SUMOylation of Small Heterodimer Partner in maintaining bile acid homeostasis. Nat. Commun. 7, 12179 (2016).

26. P. Morgan et al., Can the flow of medicines be improved? Fundamental pharmacokinetic and pharmacological principles toward improving Phase II survival. Drug Discov. Today. 17, 419–424 (2012).

27. M. Bantscheff et al., Chemoproteomics profiling of HDAC inhibitors reveals selective targeting of HDAC complexes. Nat. Biotechnol. 29, 255–265 (2011).

28. Assessment report Farydak, procedure Nr. EMEA/H/C/003725/0000, 2015.

29. N. Khan et al., Determination of the class and isoform selectivity of small-molecule histone deacetylase inhibitors. Biochem. J. 409, 581–589 (2008).

